# Cortisol dynamics and GR-dependent feedback regulation in zebrafish larvae exposed to repeated stress

**DOI:** 10.1101/2024.08.07.606983

**Authors:** Luis A. Castillo-Ramírez, Soojin Ryu, Rodrigo J. De Marco

**Author notes:** **Author for correspondence:** De Marco, Rodrigo < >. **Disclosure Statement:** The authors have nothing to disclose.

## Abstract

Zebrafish larvae show a rapid increase in cortisol in response to acute stressors, followed by a decline. While these responses are documented, both the duration of the refractory period to repeated stressors and the role of glucocorticoid receptors (GR) in specific phases of the glucocorticoid negative feedback are still being clarified. We explored these questions using water vortices as stressors, combined with GR blockage and measurements of whole-body cortisol in zebrafish larvae subjected to single and repeated stress protocols. Cortisol levels were elevated 10 minutes after stress onset and returned to baseline within 30-40 minutes, depending on the stressor strength. In response to homotypic stress, cortisol levels rose above baseline if the second stressor occurred 60 or 120 minutes after the first, but not with a 30-minute interval. This suggests a rapid cortisol-mediated feedback loop with a refractory period of at least 30 minutes. Treatment with a GR blocker delayed the return to baseline and suppressed the refractory period, indicating GR-dependent early-phase feedback regulation. These findings are consistent with mammalian models and provide a framework for further analyses of early-life cortisol responses and feedback in zebrafish larvae, ideal for non-invasive imaging and high-throughput screening.

**Summary statement:** This study investigates cortisol responses in zebrafish larvae to acute stress, revealing a rapid feedback loop and the glucocorticoid receptor’s role, offering insights for future research in stress biology.

## Introduction

Glucocorticoids (GCs) are steroid hormones crucial for managing stress and aiding bodily adaptation to challenge (de Kloet et al., 1998; McEwen, 2007). Beyond their role in stress response, they are involved in metabolism, maintaining water and electrolyte balance, immune response, growth, cardiovascular function, mood, cognitive processes, reproduction, and development. GCs are primarily produced in the adrenal cortex, alongside aldosterone and dehydro-epi-androsterone, all derived from cholesterol. Additionally, GCs are synthesized in extra-adrenal sites such as the thymus, brain, and epithelial barriers, where they exert localized effects, contributing to spatial specificity in steroid actions, independent of systemic and stress-induced variations (see Timmermans et al., 2019 for a recent review).

In both mammals and zebrafish, GCs are regulated by the hypothalamic-pituitary-adrenal (HPA) axis and the homologous hypothalamic-pituitary-interrenal (HPI) axis, respectively. The HPA axis in mammals is homologous to the HPI axis in zebrafish, making zebrafish a valuable model for stress research (Alsop and Vijayan, 2008; Alsop and Vijayan, 2009; Egan et al., 2009; Champagne and Richardson, 2013; Biran et al., 2015; Eachus et al., 2021; Tan et al., 2022; Swaminathan et al., 2023; Herget et al., 2023). Both mammals and zebrafish process stress signals through analogous centres and key neuropeptides.

Corticotropin-releasing hormone (CRH) from the nucleus preopticus (NPO), homologous to the mammalian paraventricular nucleus (PVN), triggers the release of adrenocorticotropic hormone (ACTH) (Alderman and Bernier, 2009). ACTH then stimulates the interrenal gland, the fish analogue of the adrenal gland, to release GCs like cortisol, a process known as glucocorticoid reactivity (GC_R_). GCs target the brain and periphery, facilitating stress adaptation (Munck et al., 1984; Chrousos, 1998).

A key aspect of GC regulation is the negative feedback loop (GC-NF), wherein GCs inhibit the release of CRH and ACTH to normalize GC levels after stress (Dallman and Yates, 1969; de Kloet et al., 1998; Dallman et al., 1994; Charmandari et al., 2005). This feedback mechanism is crucial for preventing excessive GC effects and maintaining neuroendocrine balance (Lightman and Conway-Campbell, 2010; Gjerstad et al., 2018; McEwen, 2007; Ulrich-Lai and Herman, 2009; Lupien et al., 2009; de Kloet and Joëls, 2023). GC-NF operates across different timescales, involves both genomic and non-genomic pathways, and relies on GCs binding to GR and mineralocorticoid receptors (MR) in the hypothalamus and pituitary to inhibit CRH and ACTH release (Kim and Iremonger, 2019; Shipston, 2022; Dallman, 2005; Evanson et al., 2010; Tasker and Herman, 2011; Joëls et al., 2013).

In zebrafish larvae, GR and MR genes are expressed as early as 2- and 4-days post-fertilization (dpf), respectively (Alsop and Vijayan, 2008). Studies indicate that disruptions in GR function and alterations in cortisol levels affect baseline cortisol and GC_R_ (Griffiths et al., 2012; Wilson et al., 2013; Castillo-Ramírez et al., 2019; Nagpal et al., 2024; De Marco et al., 2013). These findings have established the zebrafish larva as a valuable model for studying cortisol response dynamics and stress mechanisms, offering both practical and biological advantages (Eachus et al., 2021; Tan et al., 2022; Swaminathan et al., 2023). The stress response system in zebrafish is highly conserved with that of mammals, making it a powerful model for investigating early life stress and its effects on brain and behaviour. Furthermore, zebrafish embryos develop externally, enabling high-throughput experiments and controlled manipulation of stress pathways that are less accessible in mammalian models. The early expression of GR and MR provides a critical window to examine how disruptions in these receptors affect cortisol regulation during development, a key element in understanding stress-related pathologies. Exploring these dynamics in zebrafish larvae provides comparative insights into glucocorticoid feedback regulation, stress adaptation, and responses to repeated stress, enhancing our understanding alongside mammalian studies. This comparative approach supports both basic and translational research.

Despite these advances, important gaps remain in our understanding of cortisol response dynamics in zebrafish larvae. Notably, the refractory period—the interval between repeated stressor exposures that allows for cortisol elevation—and the role of GR during the early phase of GC-NF are still largely underexplored. While GR-dependent feedback mechanisms are well-established in mammals, their function and timing in zebrafish larvae need further analysis. The early phase of GC-NF occurs within minutes and involves rapid GR-mediated inhibition of hormone secretion, as shown in anterior pituitary cells. This fast, GR-driven feedback is crucial as it enables the pituitary to respond dynamically to varying glucocorticoid pulses under both stress and non-stress conditions (Shipston, 2022). Additionally, while classical genomic pathways are recognized, emerging evidence underscores the significance of non-genomic mechanisms, such as modulation of membrane excitability and calcium signalling, in early GC inhibition. These mechanisms may involve both classical and membrane-associated GR, with potential cross-talk between genomic and non-genomic pathways that demand further exploration (Kim and Iremonger, 2019). Specifying the refractory period is necessary for optimizing experimental designs aimed at studying stress recovery and adaptation. It can enhance high-throughput behavioural assays, optogenetic manipulations, and transcriptomic or proteomic analyses, ultimately contributing to a more comprehensive understanding of cortisol response dynamics in zebrafish larvae.

To address these gaps, we tested two hypotheses using controlled water vortices as stressors (Castillo-Ramírez et al., 2019), alongside GR blockage, measuring GC_R_ under single and repeated stress protocols. First, we hypothesized that encountering the same acute stressor twice would result in a diminished GC_R_, with the reduction depending on the interval between exposures. We predicted that shorter intervals between stress exposures would lead to a more significant reduction in GC_R_, aiming to approximate the length of the refractory period. Our results confirmed this. The second exposure to the same stressor diminished GC_R_, with more pronounced reductions occurring at shorter intervals. Second, we hypothesized that blocking GR during the refractory period would alter cortisol response dynamics. We predicted that larvae treated with the GR blocker Mifepristone would show a higher GC_R_ to the second stress exposure compared to controls. The results revealed that Mifepristone-treated larvae showed prolonged cortisol release but retained GC_R_, unlike controls, suggesting that GR plays a key role in the early phase of GC-NF in zebrafish larvae. These findings establish a foundation for future research that links early HPI axis activation patterns with GC regulation.

## Results

### Cortisol reactivity in response to single and repeated stress

Zebrafish larvae adapt to water vortex flows through rheotaxis, a behaviour where they orient against the current, which imposes high energy demands and can activate the HPI axis. In previous work (Castillo-Ramírez et al., 2019), we showed that controlled vortex flows at specific revolutions per minute (rpm) induce a rapid and significant rise in cortisol levels, unlike conditions without flow, establishing it as an effective stressor. Although we have not directly measured the metabolic or cardiorespiratory effects of vortex exposure, elevated cortisol levels indicate considerable energy demands and physiological stress. Vortex flows offer several advantages as a stressor. They minimize confounding variables associated with other stressors, such as changes in salt concentration or pH, which can introduce non-specific effects, and they exploit an innate, reproducible behaviour, reducing experimental variability. As vortex strength (rpm) increases, cortisol levels rise proportionally, as does a larva’s swimming effort to counteract the flow (Castillo-Ramírez et al., 2019). This quantifiable relationship between vortex intensity and GC_R_ allows precise categorization of stress levels. Combined, these elements make it well-suited for repeated stress assays. Given these advantages, we used vortex-induced forced swimming to assess the effects of repeated stress exposure and GR blockage on cortisol dynamics. First, we measured baseline and post-stress whole-body cortisol in 6 dpf larvae exposed to 3-minute vortex flows of low or high strength (see Methods). Cortisol levels were assessed at 10, 20, 30, and 40 minutes after exposure (Fig. 1A). Confirming prior findings (Castillo-Ramírez et al., 2019), we found a positive correlation between vortex strength and GC_R_ (Fig. 1B, two-way ANOVA on log-transformed data: strength: F(1,40)=78.13, *p*<0.0001; time: F(3,40)=91.99, *p*<0.0001; strength x time: F(3,40)=1.59, *p*=0.21; Bonferroni’s tests: 10’: *p*<0.0001, 20’: *p*<0.0001, 30’: *p*<0.0001, 40’: *p*=0.054; N=6 per group). Whole-body cortisol levels were elevated 10 minutes post-exposure and gradually decreased thereafter. For both low and high vortex strengths, cortisol levels returned to baseline within 30 and 40 minutes, respectively (Fig. 1C, one sample *t*-tests against a mean fold change of 1: left, 10’: t(5)=7.9, *p*=0.0005; 20’: t(5)=9.5, *p*=0.0002; 30’: t(5)=1.9, *p*=0.12; 40’: t(5)=0.8, *p*=0.44; right, 10’: t(5)=6.7, *p*=0.001; 20’: t(5)=8.7, *p*=0.0003; 30’: t(5)=7.7, *p*=0.0006; 40’: t(5)=0.25, *p*=0.81). Therefore, higher cortisol responses do not hinder a rapid return to baseline whole-body cortisol after acute stress. Cortisol decreases steadily after exposure, irrespective of peak stressor-derived levels. We then explored the link between GC_R_ and a single homotypic stress event, varying the time interval between first and second vortices at 30, 60, or 120 minutes (Fig. 1D). Larvae showed cortisol levels similar to baseline when exposed to a second vortex, regardless of its strength, 30 minutes after the initial exposure (Fig. 1E, two-way ANOVA: strength: F(1,40)=40.5, *p*<0.0001; time interval: F(3,40)=34.7, *p*<0.0001; strength x time interval: F(3,40)=4.7, *p*=0.007; Bonferroni’s tests: 1st: *p*<0.0001, 30’ after 1st: *p*>0.99, 60’ after 1st: *p*=0.01, 120’ after 1st: *p*=0.0002; N=6 per group). However, at both 60 and 120 minutes post-initial exposure, they showed a GC_R_ level correlating with the strength of the second vortex (Fig. 1F, one sample *t*-tests against a mean fold change of 1: left, 30’ after 1st: t(5)=1.40, *p*=0.22; 60’ after 1st: t(5)=3.8, *p*=0.013; 120’ after 1st: t(5)=6.3, *p*=0.002; right, 30’ after 1st: t(5)=1.44, *p*=0.21; 60’ after 1st: t(5)=6.1, *p*=0.002; 120’ after 1st: t(5)=6.4, *p*=0.001).

**Figure 1.**
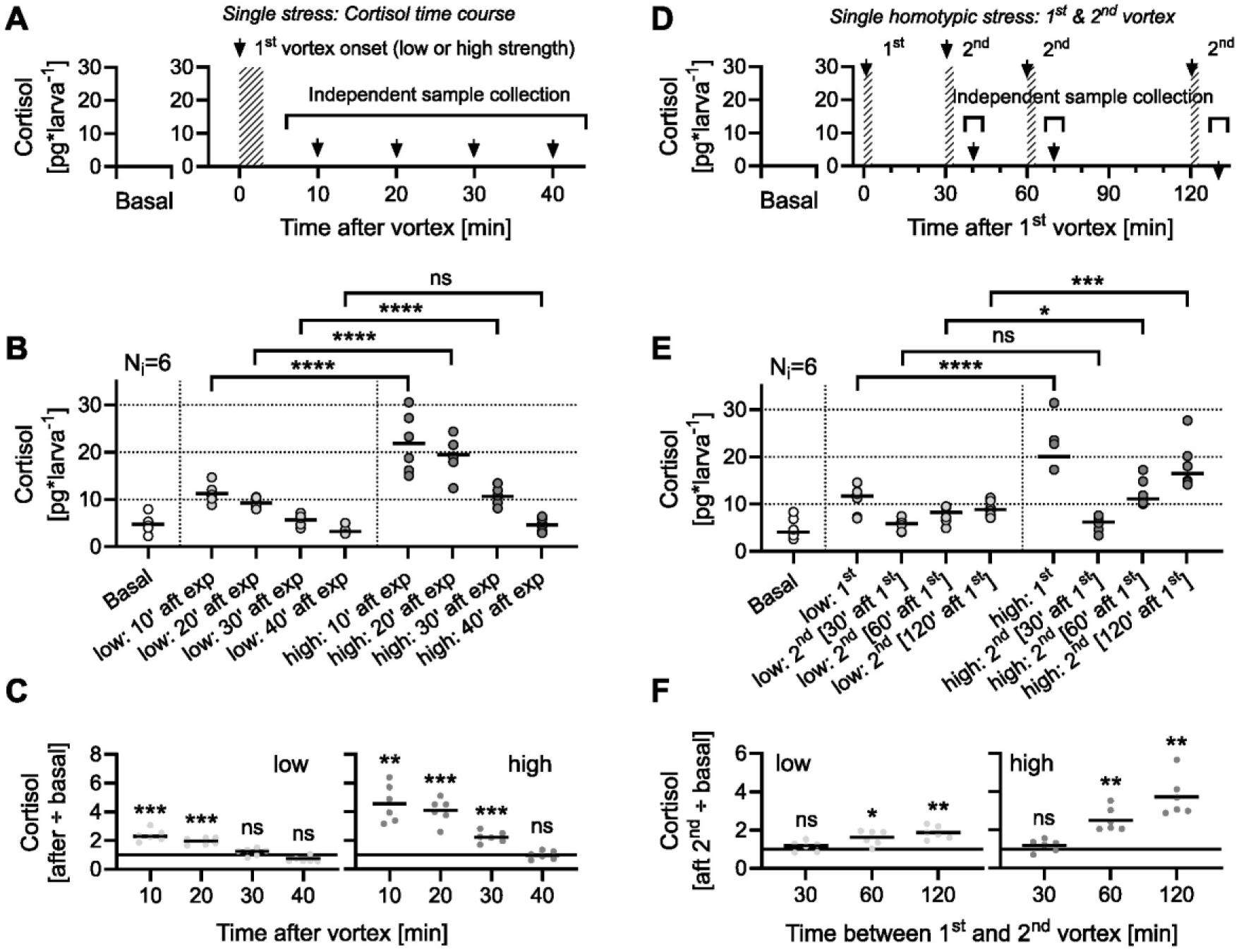
Cortisol reactivity under single and homotypic stress. (**A**) Schematic for assessing basal and post-vortex cortisol levels, measured at 10, 20, 30, and 40 minutes after exposure. (**B**) Basal and post-vortex cortisol levels. White, light grey, and dark grey represent basal cortisol and data from low and high vortex strengths, respectively, in all plots. N=6 per group. (**C**) Cortisol levels post-exposure relative to basal cortisol (data from B). (**D**) Schematic for measuring basal cortisol and GC_R_ to homotypic stress at varying intervals between two vortices (30, 60, or 120 minutes). (**E**) Basal cortisol and GC_R_ to a second vortex at varying intervals. N=6 per group. (**F**) Cortisol levels post-second exposure relative to basal cortisol (data from E). (**B**, **E**) Asterisks indicate results from Bonferroni’s tests following two-way ANOVA. (**C**, **F**) Asterisks indicate results from one sample *t*-tests against a mean fold change of 1. See Results for details on comparisons.

### Cortisol response to 30-min homotypic stress in Mifepristone-treated larvae

We then measured whole-body cortisol levels in separate groups of unstressed and stressed 6 dpf larvae treated with either DMSO or a combination of DMSO and Mifepristone, a verified zebrafish GR antagonist (Weger et al., 2012; De Marco et al., 2013). Stressed larvae experienced either a single 3-minute high-strength vortex, with cortisol levels measured at 10, 30, and 40 minutes post-initial exposure (Fig. 2A, top), or two vortices spaced 30 minutes apart (Fig. 2A, bottom). Mif-treated larvae showed similar baseline whole-body cortisol levels as controls. Additionally, while DMSO-treated larvae showed a gradual post-exposure decrease in cortisol, Mif-treated larvae retained cortisol levels above baseline at the same time point (Fig. 2B, two-way ANOVA on log-transformed data (white and black circles only): treatment: F(1,48)=65.9, *p*<0.0001; time: F(3,48)=153.3, *p*<0.0001; treatment x time: F(3,48)=24.13, *p*<0.0001; Bonferroni’s tests: basal: *p*>0.99, 10’: *p*>0.99, 30’: *p*<0.0001, 40’: *p*<0.0001; N=7 per group). As a result, DMSO-treated larvae returned to baseline cortisol levels 40 minutes post-exposure, while Mif-treated larvae did not (Fig. 2C, one sample *t*-tests against a mean fold change of 1: left, 10’: t(6)=9.0, *p*=0.0001; 30’: t(6)=4.4, *p*=0.0046; 40’: t(6)=0.2, *p*=0.85; right, 10’: t(6)=22.1, *p*<0.0001; 30’: t(6)=25.1, *p*<0.0001; 40’: t(6)=13.3, *p*<0.0001). Notably, following exposure to a second high-strength vortex 30 minutes after the initial exposure, Mif-treated larvae showed a significant cortisol increase compared to cortisol levels at the same post-initial exposure time (40 minutes), unlike DMSO-treated larvae (Fig. 2B, black and white triangles vs. circles, respectively; unpaired two-tailed t-tests: Mif-treated larvae: t(12)=5.7, *p*<0.0001; DMSO-treated larvae: t(12)=0.12, *p*=0.91). Thus, unlike untreated or DMSO-treated larvae, Mif-treated larvae showed a significant GC_R_ to homotypic stress applied during the 30-minute refractory period (Fig. 2D, one sample *t*-tests against a mean fold change of 1: untreated: t(5)=0.74, *p*=0.49; DMSO-treated: t(6)=0.17, *p*=0.87; Mif-treated: t(6)=7.2, *p*=0.0004; N per group = 7, except for the untreated group (D), where N = 6).

**Figure 2.**
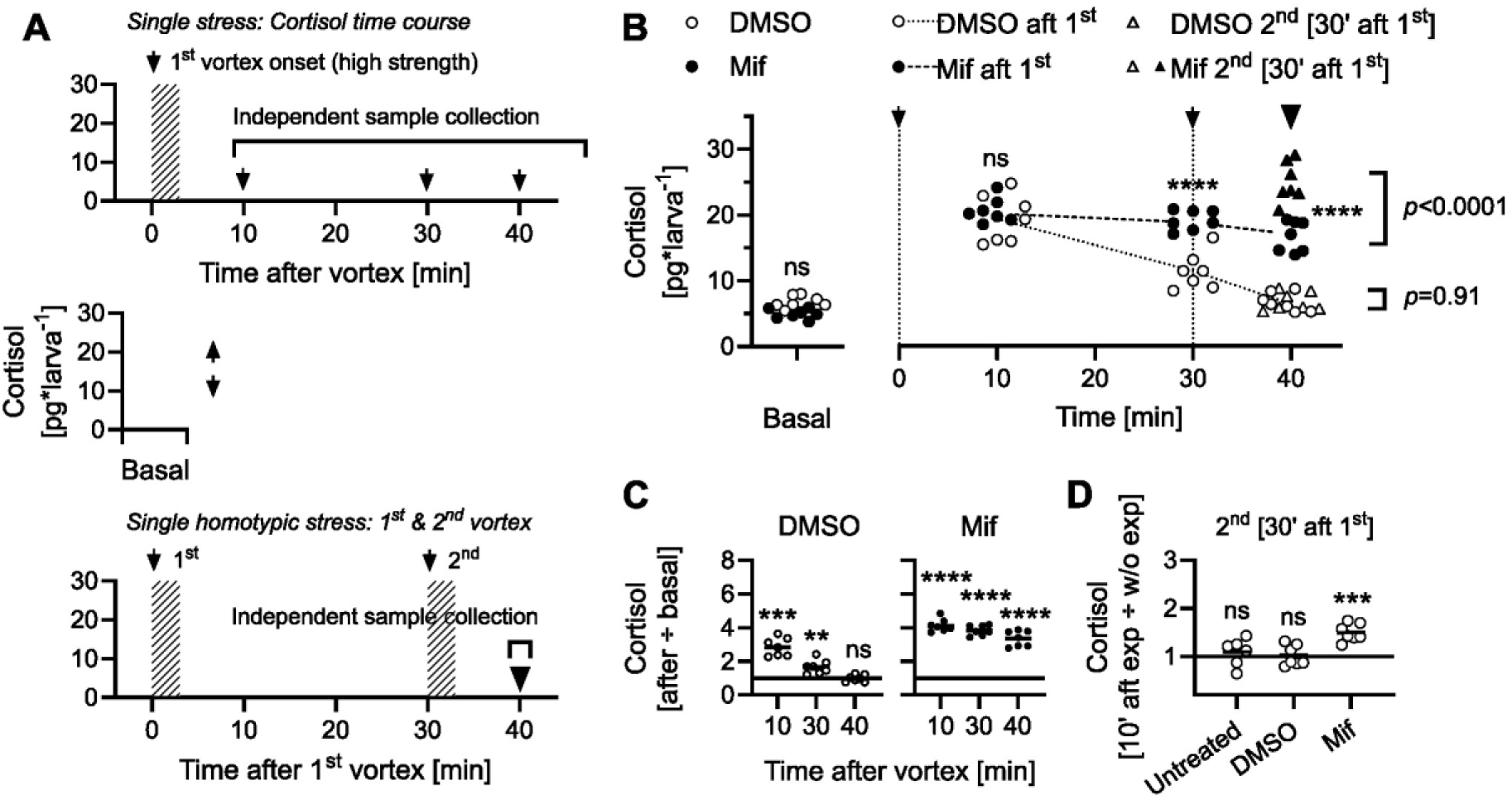
Cortisol response to 30-min homotypic stress in Mifepristone-treated larvae. (**A**) Schematic for assessing basal cortisol, post-vortex cortisol levels (measured at 10, 30, and 40 minutes post-exposure), and the GC_R_ to a second high-strength vortex (with a 30-minute interval) in larvae incubated with either DMSO or DMSO + Mifepristone (Mif). For all plots, except D, white represents DMSO group data, and black represents Mif group data. N per group = 7, except for the untreated group (D), where N = 6. (**B**) Baseline cortisol, post-first vortex levels, and cortisol in DMSO- and Mif-treated larvae after a second vortex at 30 minutes post-initial exposure, compared to independently collected cortisol levels at the same time (40 minutes). Asterisks indicate results from Bonferroni’s tests following two-way ANOVA (white and black circles only); *p* values for black and white triangles vs. circles are from unpaired two-tailed t-tests. (**C**) Cortisol levels post-first vortex relative to baseline cortisol (data from B). (**D**) Cortisol level 10 minutes after the second vortex compared to levels at the same time without the vortex (data from B, 40 minutes). (**C**, **D**) Asterisks indicate results from one sample *t*-tests against a mean fold change of 1. See Results for details on comparisons.

## Discussion

We further examined GC_R_ dynamics and cortisol feedback in stressed zebrafish larvae, focusing on the refractory period in GC_R_ to repeated stress and the role of GR in the early phase of GC-NF. We observed elevated cortisol levels 10 minutes after the onset of stress, with levels returning to baseline 30-40 minutes later, depending on the strength of the stressor. Our measurements of cortisol at 5, 10, and 15 minutes following vortex exposure also revealed a peak at 10 minutes post-vortex onset (data not shown). This rapid increase in cortisol is consistent with previous findings, which report elevated whole-body cortisol levels around 10 minutes after stress in zebrafish larvae exposed to brief vortices, mild electric shocks, and air exposure (Faught and Vijayan, 2022; Hare et al., 2021; Steenbergen et al., 2012). In response to a repeated stressor, cortisol levels rose above baseline only when the second stressor occurred 60 or 120 minutes after the first, but not with a 30-minute interval. These results indicate a rapid cortisol-mediated feedback loop with a refractory period of at least 30 minutes. The return to baseline cortisol levels was delayed, and the refractory period was suppressed when larvae were treated with the GR antagonist Mifepristone, indicating GR-dependent feedback. These findings are crucial for validating the relevance of zebrafish as a stress research model.

Previous evidence links GR to GC-NF in zebrafish larvae. Both GR and MR contribute to early cortisol balance (Alsop and Vijayan, 2009; Faught and Vijayan, 2018), and GR-deficient larvae fail to show typical cortisol responses to stress, with cortisol levels remaining elevated, indicative of impaired negative feedback mechanisms. These mutants show increased HPI axis activity, reduced cortisol suppression by dexamethasone, and elevated CRH and ACTH levels (Griffiths et al., 2013; Ziv et al., 2013; Fonseka et al., 2016). Cortisol responses to acute stressors, such as light pulses, show temporal patterns similar to our findings, with an initial rise followed by a return to baseline (De Marco et al., 2013; De Marco et al., 2014; De Marco et al., 2016). Notably, repeated light pulses delivered at 30-minute intervals do not lead to additional cortisol elevation after the first pulse, suggesting a refractory period (De Marco et al., 2013). This pattern is consistent with our results, where a second stressor applied after a 30-minute interval also failed to elicit a significant cortisol rise, reinforcing the interpretation of a 30-minute refractory period in zebrafish larvae. Unlike knockout studies, which are limited in assessing the dynamics of processes dependent on the targeted element, using a GR blocker allows for measurement of GR inhibition’s effects on GC_R_ dynamics, revealing the role of GR in cortisol feedback regulation. Pre-incubation with the GR blocker Mifepristone elevated cortisol levels for both initial and subsequent light pulses (De Marco et al., 2013), similar to our observation that Mifepristone-treated larvae showed prolonged cortisol release after repeated vortex stress. This correspondence between light pulse and vortex-induced stress responses supports the role of GR in modulating GC_R_ dynamics and strengthens our interpretation of the 30-minute refractory period observed in repeated stress assays.

Our findings confirmed that repeated exposure to the same stressor reduces GC_R_, with shorter intervals between exposures yielding greater reductions, and that blocking GR during the refractory period prolongs cortisol release while retaining GC_R_. Confirming both predictions is important because the reduction in GC_R_ to closely spaced stressors could involve neural mechanisms independent of glucocorticoids, similar to what has been observed in mice. In mice, stress familiarity reduces CRH neuron responses without GC-NF involvement, potentially limiting HPA axis activation (Kim et al., 2019). Although this possibility requires further exploration in zebrafish, our results collectively suggest that early-phase GR-mediated feedback reduces GC_R_ in zebrafish larvae when exposed to repeated homotypic stress within 30 minutes.

Mifepristone is a well-established GR antagonist, and the dose and incubation period we used follow verified protocols for zebrafish larvae (Weger et al., 2012; De Marco et al., 2013). However, Mifepristone may have off-target effects, such as antagonizing progesterone receptors and potentially interacting with other receptors at higher concentrations. These non-target effects should be considered when interpreting some of our results. Future studies should explore alternative GR antagonists and evaluate GR-responsive gene expression to gain more insights into GC_R_ dynamics with GR inhibition. Additionally, exploring how GR antagonists affect baseline cortisol levels in relation to age, dosage, and incubation time will be necessary for a comprehensive understanding. In zebrafish larvae, MR signalling also plays a role in regulating the HPI axis. MR mutants do not show differences in baseline or maximum post-stress cortisol levels, but they show a delayed cortisol rise and prolonged elevation beyond 30 minutes post-stress compared to controls (Faught and Vijayan, 2018). Further research is needed to clarify the distinct and combined roles of GR and MR in early GC-NF. Also, addressing age-dependent and long-term effects of GR blockade on stress physiology, as well as replicating these experiments with heterotypic stressors (e.g., Herget et al., 2023), will advance our understanding of acute stress responses in zebrafish larvae.

Data on the relationship between GR, GC-NF, and GC_R_ during repeated stress in rodents is limited. Past rodent studies examined ACTH and corticosterone secretion after restraint stress (De Souza and Van Loon, 1982). After a single stress event, ACTH peaked quickly and normalized within 30 minutes, while corticosterone peaked at 15-30 minutes and returned to baseline within 60-90 minutes. These responses were consistent after repeated stress every 90 minutes, but closer stress events reduced the corticosterone response, indicating a narrower adrenocortical window. Compelling evidence indicates that GR activation regulates stress-induced HPA axis activity (Ratka et al., 1989; Sapolsky et al., 2000). Local GR knockdown in the PVN increased stress-induced ACTH and corticosterone (Solomon et al., 2015), and genetic manipulations of GR elevated stress-induced GCs and disrupted GC-NF (Laryea et al., 2013). Thus, PVN GR mediates early-phase feedback, controlling the duration and intensity of GC secretion post-stress. The complex nature of GC-NF during acute stress is underscored by the fact that GR activation in the brain supports arousal, cognitive performance, and modulates immune and inflammatory responses to meet metabolic demands (de Kloet, 2023). Our study explores how GR-mediated feedback impacts GC_R_ dynamics in zebrafish larvae, a less commonly studied model organism in this context. Like rodents, zebrafish larvae show a rapid cortisol response to initial stress followed by a refractory period with repeated stress. Thus, our findings help inform experimental design and facilitate comparisons between zebrafish and mammalian models. This is particularly relevant in the context of development, as larvae offer an opportunity to explore early patterns of HPI axis activity.

Our data not only reflect GC_R_ dynamics seen in conventional mammalian models but also demonstrate the potential of the high-throughput forced swim procedure for addressing repeated stress in zebrafish larvae. This is in line with the 3Rs principle, as zebrafish larvae are increasingly recognized for their value in stress research (e.g., Eachus et al., 2021; Swaminathan et al., 2023; De Marco et al., 2016; vom Berg-Maurer et al., 2016; de Abreu et al., 2021). Collectively, the results provide a more detailed framework for exploring the link between early life stress and HPI axis programming, as well as for dissecting GC regulation in the NPO and pituitary. With the small size and transparency of zebrafish larvae, future studies can image stress-related activity patterns in both regions. Understanding how the refractory period and GR blocker sensitivity influence these patterns can enhance our grasp of cortisol response timing. This knowledge may help identify mechanisms by which the developing HPI axis adapts or fails to adapt to homotypic stress.

## Methods

### Zebrafish husbandry, handling, and experimental unit

Zebrafish breeding and maintenance were conducted under standard conditions (Westerfield, 2000). Groups of thirty wild-type eggs (cross of AB and TL strains, AB/TL) were collected in the morning, and the embryos were raised on a 12:12 light/dark cycle at 28°C in 35 mm Petri dishes with 5 ml of E2 medium. The modified Embryo-Medium2 (0.5x E2, 1 L) consisted of 5 mM NaCl, 0.25 mM KCl, 0.5 mM MgSO_4_ × 7 H_2_O, 0.15 mM KH_2_PO_4_, 0.05 mM Na_2_HPO_4_, 0.5 mM CaCl_2_, and 0.71 mM NaHCO_3_. This formulation provides a balanced ionic environment suitable for zebrafish embryo development. At 3 dpf, the E2 medium was renewed, and chorions and debris were removed from the dishes. Experiments were performed with 6 dpf larvae. In all experiments, each experimental unit (replicate) comprised a group of thirty larvae, maintained in a 35 mm Petri dish. All dishes were kept under identical conditions in the incubator on top of the stirrer plate (see below) to ensure no perturbation. Zebrafish experimental procedures were conducted according to the guidelines of the German animal welfare law and were approved by the local government (Regierungspräsidium Karlsruhe; G-29/12).

### Water vortex flows

We used water vortices in a high-throughput fashion to induce rheotaxis and cortisol increase (Castillo-Ramírez et al., 2019). The groups of thirty larvae in 35 mm Petri dishes with 5 ml of E2 medium (experimental units) were exposed to controlled vortices generated by the spinning movements of small encapsulated (coated) magnetic stir bars (6 x 3mm, Fischerbrand, #11888882, Fisher scientific, Leicestershire, UK) inside the dishes. The Petri dishes, each with a single stir bar, were always positioned on magnetic stirrer plates (Variomag, Poly 15; Thermo Fisher Scientific, Leicestershire, UK) and kept at 28°C inside an incubator (RuMed 3101, Rubarth Apparate GmbH, Laatzen, Germany). Larvae were exposed to vortex flows only when induced by the magnetic field inversions of the stirrer plate for 3-minute periods, at 130 (low strength) or 330 (high strength) revolutions per minute (rpm). The magnetic field inversions of the stirrer plate, *per se*, do not alter the level of whole-body cortisol in larval zebrafish, and we used a maximum of 330 rpm to avoid potential ceiling effects caused by the maximum levels of vortex-dependent cortisol increase, as previously established (Castillo-Ramírez et al., 2019). Larvae were exposed to the 3-min vortex either once or twice, with the second exposure occurring 30, 60, or 120 minutes after the first exposure, depending on the experimental series. These time intervals were chosen to balance accurate estimation of the cortisol refractory period with the 3Rs principle, minimizing the number of subjects. The 30, 60, and 120-minute intervals effectively capture cortisol recovery dynamics following an initial stressor. They provide a range broad enough to account for variations in stress response timing, while remaining precise enough to determine when cortisol levels return to baseline or become responsive to a subsequent stressor. This approach ensured experimental rigor while adhering to ethical guidelines for subject use. The larvae in groups of thirty were subsequently used for cortisol detection (see below). Basal levels of cortisol were measured at the time corresponding to the beginning of the experiment, before the first exposure to vortex flows.

### Whole-body cortisol

The procedures for cortisol measurements and the homemade ELISA were as previously described (Yeh et al., 2013). Validation of the homemade ELISA was conducted through intra-assay and inter-assay precision, recovery rates, and cross-reactivity assessments. Intra-assay precision was confirmed with coefficients of variation (CVs) of 10.1%, 13.7%, and 9.0% for low (1.6 ng/mL), medium (5.6 ng/mL), and high (17.7 ng/mL) cortisol concentrations, respectively. Inter-assay precision, measured over five days, yielded CVs of 10.5%, 18.5%, and 7.6% for low (1.6 ng/mL), medium (5.7 ng/mL), and high (20.7 ng/mL) concentrations. Recovery rates were high, ranging from 85% to 100% for larval samples and up to 145% for adult samples. Matrix effects were corrected using a recovery function to ensure accurate measurements. Additionally, comparison with a commercial kit showed no significant differences, and baseline cortisol levels were consistent across batches and sessions (one-way ANOVA, F(4,20)=0.035, *p*=0.99), confirming reliability of the assay. Following vortex exposure, groups of 30 larvae (replicates) were immobilized in ice water, frozen in an ethanol/dry-ice bath, and stored at −20°C for subsequent cortisol extraction and detection between 10:30 and 11:30 hours.

### Mifepristone incubation

Before exposure to the vortex, larvae were incubated for 2 hours in 1 µM Mifepristone (RU486, Sigma-Aldrich) dissolved in E2 Medium with 0.1% DMSO. These parameters were based on established protocols (Weger et al., 2012; De Marco et al., 2013) to ensure consistency with verified procedures and facilitate direct comparisons with existing literature.

### General design and statistical analysis

As stated above, measurements were conducted on distinct groups of thirty larvae (replicates). Larval density per well was kept constant. For each measurement, all thirty larvae within a well were used. Each replicate was fully independent of the others. In all experiments, treatments were randomly assigned to replicates, and blinding was implemented. An initial experimenter conducted the treatments, collected, and labelled the samples. A second experimenter then performed the measurements on the labelled samples, assigning new labels. The first experimenter subsequently quantified cortisol using these newly encoded samples. Our sample sizes are consistent with those commonly used in the field and align with previous publications (Castillo-Ramírez et al., 2019; De Marco et al., 2013; De Marco et al., 2016; Yeh et al., 2013; Herget et al., 2023; vom Berg-Maurer et al., 2016). They are based on prior work that established acceptable coefficients of variation for cortisol measurements while minimizing subject use in accordance with ethical guidelines. Data were tested for normality and homoscedasticity using the Shapiro-Wilk and KS normality tests and the Brown–Forsythe and Bartlett’s tests, respectively. We employed two-way ANOVAs followed by Bonferroni’s post-hoc tests for multiple comparisons. In cases where the data did not meet the assumption of homoscedasticity, we applied log transformations to normalize variance. Unpaired two-tailed t-tests were used for comparing independent groups, and one-sample t-tests were performed when comparing data to a hypothetical mean (i.e., a fold change of 1). For one-sample t-tests, Bonferroni corrections were applied by dividing the significance level (alpha) by the number of comparisons to control for multiple testing. No data points or samples were excluded from the analysis. Statistical analyses were performed using MS-Excel (Microsoft Corp; Redmond, WA, USA) and Prism 10.2.0 (Graphpad Software Inc, San Diego, CA, USA).

## Acknowledgments

We thank Karl J. Iremonger for providing useful comments on a previous draft, R. Singer and A. Schoell for their expertise in fish care, and R. Rödel for technical support. Also, we would like to thank the reviewers for their comments, which improved the quality of this manuscript.

## Data availability

The experimental datasets presented in this study are available upon request from the authors.

## Author Contributions

Luis A. Castillo-Ramírez, Conceptualization, Data curation, Formal analysis, Investigation, Validation, Methodology, Writing - original draft; Soojin Ryu, Conceptualization, Resources, Supervision, Funding acquisition, Methodology, Project administration, Writing – review; Rodrigo J. De Marco, Conceptualization, Resources, Supervision, Funding acquisition, Investigation, Validation, Methodology, Data curation, Formal analysis, Visualization, Project administration, Writing - original draft, Writing - review and editing.

## References

Alderman, S.L. and Bernier, N.J. (2009). Ontogeny of the corticotropin-releasing factor system in zebrafish. Gen. Comp. Endocrinol. 164(1),61–9. doi: 10.1016/j.ygcen.2009.04.007.

Alsop, D. and Vijayan, M.M. (2008). Development of the corticosteroid stress axis and receptor expression in zebrafish. Am. J. Physiol. Regul. Integr. Comp. Physiol. 294(3),R711–9. doi10.1152/ajpregu.00671.2007

Alsop, D. and Vijayan, M.M. (2009). Molecular programming of the corticosteroid stress axis during zebrafish development. Comp. Biochem. Physiol. A. Mol. Integr. Physiol. 153(1),49–54. doi10.1016/j.cbpa.2008.12.008

Biran, J., Tahor, M., Wircer, E. and Levkowitz, G. (2015). Role of developmental factors in hypothalamic function. Front. Neuroanat. 9,47. doi: 10.3389/fnana.2015.00047.

Castillo-Ramírez, L.A., Ryu, S., and De Marco, R.J. (2019). Active behaviour during early development shapes glucocorticoid reactivity. Sci. Rep. 9(1),12796. doi: 10.1038/s41598-019-49388-3.

Champagne, D.L., and Richardson, M.K. (2013). Patterns of cortisol reactivity to stress and the development of coping style in zebrafish *Danio rerio*. J. Neuroendocrinol. 25(9),750–61. doi: 10.1111/jne.12053.

Charmandari, E., Tsigos, C., and Chrousos, G. (2005). Endocrinology of the stress response. Annu. Rev. Physiol. 67,259–84. doi: 10.1146/annurev.physiol.67.040403.120816.

Chrousos, G.P. (1998). Stressors, stress, and neuroendocrine integration of the adaptive response. The 1997 Hans Selye Memorial Lecture. Ann. N. Y. Acad. Sci. 851,311–35. doi: 10.1111/j.1749-6632.1998.tb09006.x.

Dallman, M.F. (2005). Fast glucocorticoid actions on brain: Back to the future. Front. Neuroendocrinol. 26(3-4),103–8. doi: 10.1016/j.yfrne.2005.08.001.

Dallman, M.F., and Yates, F.E. (1969). Dynamic asymmetries in the corticosteroid feedback path and distribution-metabolism-binding elements of the adrenocortical system. Ann. N. Y. Acad. Sci. 156(2),696–721. doi: 10.1111/j.1749-6632.1969.tb14008.x.

Dallman, M.F., Akana, S.F., Levin, N., Walker, C.D., Bradbury, M.J., Suemaru, S., and Scribner, K.S. (1994). Corticosteroids and the control of function in the hypothalamo-pituitary-adrenal (HPA) axis. Ann. N. Y. Acad. Sci. 746,22–31. doi: 10.1111/j.1749-6632.1994.tb39206.x.

de Abreu, M.S., Demin, K.A., Giacomini, A.C.V.V., Amstislavskaya, T.G., Strekalova, T., Maslov, G.O., Kositsin, Y., Petersen, E.V., and Kalueff, A.V. (2021). Understanding how stress responses and stress-related behaviors have evolved in zebrafish and mammals. Neurobiol. Stress. 15,100405. doi: 10.1016/j.ynstr.2021.100405.

de Kloet, E.R. (2023). Glucocorticoid feedback paradox: a homage to Mary Dallman. Stress 26(1):2247090. doi: 10.1080/10253890.2023.2247090.

de Kloet, E.R., Vreugdenhil, E., Oitzl, M.S. and Joëls, M. (1998) Brain corticosteroid receptor balance in health and disease. Endocr. Rev. 19(3):269–301. doi: 10.1210/edrv.19.3.0331.

de Kloet, E.R., and Joëls, M. (2023). The cortisol switch between vulnerability and resilience. Mol. Psychiatry 29(1),20–34. doi: 10.1038/s41380-022-01934-8.

De Marco, R.J., Groneberg, A.H., Yeh, C.M., Castillo Ramírez, L.A., and Ryu, S. (2013). Optogenetic elevation of endogenous glucocorticoid level in larval zebrafish. Front. Neural Circuits 7,82. doi: 10.3389/fncir.2013.00082.

De Marco, R.J., Groneberg, A.H., Yeh, C.M., Treviño, M., and Ryu, S. (2014). The behavior of larval zebrafish reveals stressor-mediated anorexia during early vertebrate development. Front. Behav. Neurosci. 8:367. doi: 10.3389/fnbeh.2014.00367.

De Marco, R.J., Thiemann, T., Groneberg, A.H., Herget, U., and Ryu, S. (2016). Optogenetically enhanced pituitary corticotroph cell activity post-stress onset causes rapid organizing effects on behaviour. Nat. Commun. 7,12620. doi: 10.1038/ncomms12620

De Souza, E.B., and Van Loon, G.R. (1982). Stress-induced inhibition of the plasma corticosterone response to a subsequent stress in rats: a nonadrenocorticotropin-mediated mechanism. Endocrinology 110(1):23–33. doi: 10.1210/endo-110-1-23.

Eachus, H., Choi, M.K., and Ryu, S. (2021). The Effects of Early Life Stress on the Brain and Behaviour: Insights From Zebrafish Models. Front. Cell Dev. Biol. 9,657591. doi: 10.3389/fcell.2021.657591.

Egan, R.J., Bergner, C.L., Hart, P.C., Cachat, J.M., Canavello, P.R., Elegante, M.F., Elkhayat, S.I., Bartels, B.K., Tien, A.K., Tien, D.H., et al. (2009). Understanding behavioral and physiological phenotypes of stress and anxiety in zebrafish. Behav. Brain Res. 205(1),38–44. doi: 10.1016/j.bbr.2009.06.022.

Evanson, N.K., Tasker, J.G., Hill, M.N., Hillard, C.J., and Herman, J.P. (2010). Fast feedback inhibition of the HPA axis by glucocorticoids is mediated by endocannabinoid signaling. Endocrinology 151(10):4811–9. doi: 10.1210/en.2010-0285.

Faught, E., and Vijayan, M.M. (2018). The mineralocorticoid receptor is essential for stress axis regulation in zebrafish larvae. Sci. Rep. 8(1):18081. doi: 10.1038/s41598-018-36681-w.

Faught, E., and Vijayan, M.M. (2022). Coordinated action of corticotropin-releasing hormone and cortisol shapes the acute stress-induced behavioural response in zebrafish. Neuroendocrinology. 112(1):74–87. doi: 10.1159/000514778.

Fonseka, T.M., Wen, X.Y., Foster, J.A., and Kennedy, S.H. (2016). Zebrafish models of major depressive disorders. J. Neurosci. Res. 94(1):3–14. doi: 10.1002/jnr.23639.

Gjerstad, J.K., Lightman, S.L., and Spiga, F. (2018). Role of glucocorticoid negative feedback in the regulation of HPA axis pulsatility. Stress 21(5):403–416. doi: 10.1080/10253890.2018.1470238.

Griffiths, B.B., Schoonheim, P.J., Ziv, L., Voelker, L., Baier, H., and Gahtan, E. (2012) A zebrafish model of glucocorticoid resistance shows serotonergic modulation of the stress response. Front. Behav. Neurosci. 6:68. doi: 10.3389/fnbeh.2012.00068.

Hare, A.J., Zimmer, A.M., LePabic, R., Morgan, A.L., and Gilmour, K.M. (2021). Early-life stress influences ion balance in developing zebrafish (*Danio rerio*). J. Comp. Physiol. B. 191(1):69–84. doi: 10.1007/s00360-020-01319-9.

Herget, U., Ryu, S. and De Marco, R.J. (2023) Altered glucocorticoid reactivity and behavioral phenotype in rx3-/- larval zebrafish. Front. Endocrinol. 6;14:1187327. doi: 10.3389/fendo.2023.1187327.

Herget, U., Wolf, A., Wullimann, M.F., and Ryu, S. (2014). Molecular neuroanatomy and chemoarchitecture of the neurosecretory preoptic-hypothalamic area in zebrafish larvae. J. Comp. Neurol. 522(7),1542–1564. doi: 10.1002/cne.23480.

Joëls, M., Pasricha, N., Karst, H. (2013). The interplay between rapid and slow corticosteroid actions in brain. Eur. J. Pharmacol. 719(1-3):44–52. doi: 10.1016/j.ejphar.2013.07.015.

Kim, J.S., and Iremonger, K.J. (2019). Temporally Tuned Corticosteroid Feedback Regulation of the Stress Axis. Trends Endocrinol. Metab. 30(11):783–792. doi: 10.1016/j.tem.2019.07.005.

Kim, J.S., Han, S.Y., and Iremonger, K.J. (2019). Stress experience and hormone feedback tune distinct components of hypothalamic CRH neuron activity. Nat. Commun. 10(1):5696. doi: 10.1038/s41467-019-13639-8.

Laryea, G., Schütz, G., and Muglia, L.J. (2013). Disrupting hypothalamic glucocorticoid receptors causes HPA axis hyperactivity and excess adiposity. Mol. Endocrinol. 27(10):1655–65. doi: 10.1210/me.2013-1187.

Lightman, S.L., and Conway-Campbell, B.L. (2010). The crucial role of pulsatile activity of the HPA axis for continuous dynamic equilibration. Nat. Rev. Neurosci. 11(10):710–8. doi: 10.1038/nrn2914.

Lupien, S.J., McEwen, B.S., Gunnar, M.R., and Heim, C. (2009). Effects of stress throughout the lifespan on the brain, behaviour and cognition. Nat. Rev. Neurosci. 10(6):434–45. doi: 10.1038/nrn2639.

McEwen, B.S. (2007). Physiology and neurobiology of stress and adaptation: central role of the brain. Physiol. Rev. 87(3),873–904. doi: 10.1152/physrev.00041.2006.

Munck, A., Guyre, P.M., and Holbrook, N.J. (1984). Physiological functions of glucocorticoids in stress and their relation to pharmacological actions. Endocr. Rev. 5(1),25–44. doi: 10.1210/edrv-5-1-25.

Nagpal, J., Eachus, H., Lityagina, O., and Ryu, S. (2024). Optogenetic induction of chronic glucocorticoid exposure in early-life leads to blunted stress-response in larval zebrafish. Eur. J. Neurosci. 59(11),3134–3146. doi: 10.1111/ejn.16301.

Ratka, A., Sutanto, W., Bloemers, M., and de Kloet, E.R. (1989). On the role of brain mineralocorticoid (type I) and glucocorticoid (type II) receptors in neuroendocrine regulation. Neuroendocrinology 50(2):117–23. doi: 10.1159/000125210.

Sapolsky, R.M., Romero, L.M., and Munck, A.U. (2000). How do glucocorticoids influence stress responses? Integrating permissive, suppressive, stimulatory, and preparative actions. Endocr. Rev. 21(1):55–89. doi: 10.1210/edrv.21.1.0389.

Shipston, M.J. (2022). Glucocorticoid action in the anterior pituitary gland: Insights from corticotroph physiology. Curr. Opin. Endocr. Metab. Res. 25,100358. doi: 10.1016/j.coemr.2022.100358.

Solomon, M.B., Loftspring, M., de Kloet, A.D., Ghosal, S., Jankord, R., Flak, J.N., Wulsin, A.C., Krause, E.G., Zhang, R., Rice, T., McKlveen, J., Myers, B., Tasker, J.G., and Herman, J.P. (2015). Neuroendocrine Function After Hypothalamic Depletion of Glucocorticoid Receptors in Male and Female Mice. Endocrinology 156(8):2843–53. doi: 10.1210/en.2015-1276.

Steenbergen, P.J., Metz, J.R., Flik, G., Richardson, M.K., and Champagne, D.L. (2012). Methods to quantify basal and stress-induced cortisol response in larval zebrafish. Neuromethods. 66, 121–141. 10.1007/978-1-61779-597-8_9

Swaminathan, A., Gliksberg, M., Anbalagan, S., Wigoda, N., and Levkowitz, G. (2023). Stress resilience is established during development and is regulated by complement factors. Cell Rep. 42(1),111973. doi: 10.1016/j.celrep.2022.111973.

Tan, J.X.M., Ang, R.J.W., and Wee, C.L. (2022). Larval zebrafish as a model for mechanistic discovery in mental health. Front. Mol. Neurosci. 15:900213. doi: 10.3389/fnmol.2022.900213.

Tasker, J.G., and Herman, J.P. (2011). Mechanisms of rapid glucocorticoid feedback inhibition of the hypothalamic-pituitary-adrenal axis. Stress 14(4):398–406. doi: 10.3109/10253890.2011.586446.

Timmermans, S., Souffriau, J., and Libert, C. (2019). A General Introduction to Glucocorticoid Biology. Front. Immunol. 10:1545. doi: 10.3389/fimmu.2019.01545.

Ulrich-Lai, Y.M., and Herman, J.P. (2009). Neural regulation of endocrine and autonomic stress responses. Nat. Re.v Neurosci. 10(6):397–409. doi: 10.1038/nrn2647.

vom Berg-Maurer, C.M., Trivedi, C.A., Bollmann, J.H., De Marco, R.J., and Ryu, S. (2016). The Severity of Acute Stress Is Represented by Increased Synchronous Activity and Recruitment of Hypothalamic CRH Neurons. J. Neurosci. 36(11):3350–62. doi: 10.1523/JNEUROSCI.3390-15.2016.

Weger, B.D., Weger, M., Nusser, M., Brenner-Weiss, G., and Dickmeis, T. (2012). A chemical screening system for glucocorticoid stress hormone signaling in an intact vertebrate. ACS Chem. Biol. 7(7):1178–83. doi: 10.1021/cb3000474.

Westerfield, M. (2000). The Zebrafish Book. A Guide for the Laboratory Use of Zebrafish (Danio rerio). 4th ed. Eugene: Univ. of Oregon Press.

Wilson, K.S., Matrone, G., Livingstone, D.E., Al-Dujaili, E.A., Mullins, J.J., Tucker, C.S., Hadoke, P.W., Kenyon, C.J., and Denvir, M.A. (2013) Physiological roles of glucocorticoids during early embryonic development of the zebrafish (*Danio rerio*). J. Physiol. 591(24):6209–20. doi: 10.1113/jphysiol.2013.256826.

Yeh, C.M., Glöck, M., and Ryu, S. (2013). An optimized whole-body cortisol quantification method for assessing stress levels in larval zebrafish. PLoS One 8(11),e79406. doi: 10.1371/journal.pone.0079406.

Ziv, L., Muto, A., Schoonheim, P.J., Meijsing, S.H., Strasser, D., Ingraham, H.A., Schaaf, M.J., Yamamoto, K.R., and Baier, H. (2013). An affective disorder in zebrafish with mutation of the glucocorticoid receptor. Mol. Psychiatry 18(6),681–91. doi: 10.1038/mp.2012.64.

